# The Metabolic Capability and Phylogenetic Diversity of Mono Lake During a Bloom of the Eukaryotic Phototroph *Picocystis* strain ML

**DOI:** 10.1101/334144

**Authors:** Blake W. Stamps, Heather S. Nunn, Victoria A. Petryshyn, Ronald S. Oremland, Laurence G. Miller, Michael R. Rosen, Kohen W. Bauer, Katharine J. Thompson, Elise M. Tookmanian, Anna R. Waldeck, Sean J. Lloyd, Hope A. Johnson, Bradley S. Stevenson, William M. Berelson, Frank A. Corsetti, John R. Spear

**Affiliations:** Department of Civil and Environmental Engineering, Colorado School of Mines, Golden, Colorado, USA; Department of Microbiology and Plant Biology, University of Oklahoma, Norman, Oklahoma, USA; Environmental Studies Program, University of Southern California, Los Angeles, California, USA; Department of Earth Sciences, University of Southern California, Los Angeles, California, USA; United States Geological Survey, Menlo Park, California, USA; United States Geological Survey, Carson City, Nevada, USA; Department of Earth, Ocean and Atmospheric Sciences, University of British Columbia, Vancouver, British Columbia, Canada.; Division of Chemistry and Chemical Engineering, California Institute of Technology, Pasadena, California, USA; Department of Earth and Planetary Sciences, Harvard University, Cambridge, Massachusetts, USA; Department of Geological Sciences, California State University Fullerton, Fullerton, California, USA; Department of Biological Science, California State University Fullerton, Fullerton, California, USA

## Abstract

Algal blooms in lakes are often associated with anthropogenic eutrophication; however, they can occur naturally. In Spring of 2016 Mono Lake, a hyperalkaline lake in California, was near the height of a rare bloom of the algae Picocystis strain ML and at the apex of a multi-year long drought. These conditions presented a unique sampling opportunity to investigate microbiological dynamics during an intense natural bloom. We conducted a comprehensive molecular analysis along a depth transect near the center of the lake from surface to 25 m depth during June 2016. Across sampled depths, rRNA gene sequencing revealed that Picocystis associated chloroplast were found at 40-50 % relative abundance, greater than values recorded previously. Despite the presence of the photosynthetic oxygenic algal genus Picocystis, oxygen declined below detectible limits below 15 m depth, corresponding with an increase in microorganisms known to be anaerobic. In contrast to previously sampled years, metagenomic and metatranscriptomic data suggested a loss of sulfate reducing microorganisms throughout the lake’s water column. Gene transcripts associated with Photosystem I and II were expressed at both 2 m and 25 m, suggesting that limited oxygen production may occur at extremely low light levels at depth within the lake. Oxygenic photosynthesis under low light conditions, in the absence of potential grazing by the brine shrimp Artemia, may allow for a cryptic redox cycle to occur in an otherwise anoxic setting at depth in the lake with the following effects: enhanced productivity, reduced grazing pressure on Picocystis, and an exacerbation of bloom.

**IMPORTANCE:** Mono Lake, California provides habitat to a unique ecological community that is heavily stressed due to recent human water diversions and a period of extended drought. To date, no baseline information exists about Mono Lake to understand how the microbial community responds to drought, bloom, and what genetic functions are lost in the water column. While previously identified anaerobic members of the microbial community disappear from the water column during drought and bloom, sediment samples suggest these microorganisms seek refuge at lake bottom or in the subsurface. Thus, the sediments may represent a type of seed bank which could restore the microbial community as a bloom subsides. Our work also sheds light on the activity of the halotolerant algae Picocystis strain ML during a bloom at Mono Lake, its ability to potentially produce oxygen via photosynthesis even under extreme low-light conditions, and how the remainder of the microbial community responds.

## Introduction

Mono Lake is a large hypersaline alkaline lake with a maximum depth of ≈ 50 m in the Mono Basin near the eastern foothills of the Sierra Nevada Mountains, California (Figure 1). It formed from the remnant of Paleolake Russell (a Pleistocene glacial lake) and has existed as a closed basin for at least 50,000 years (1). Diversion of tributary streams to Mono Lake by the city of Los Angeles began in 1941, and resulted in a drop of over 13 m in lake level by 1978 (2) with a corresponding increase in water salinity from 48 g/L to 81 g/L by the 1990s (3) and a current alkalinity of 30,400 ppm HCO_3_^-^ (4). The steep decline in lake level also resulted in increasing concentrations of other solutes (including arsenic), resulting in unusual lake geochemistry and a the absence of large macrofauna (e.g., fish) (5). Mono Lake is home to a photosynthetic eukaryotic algae, *Picocystis* (6) that is the primary food source of a brine shrimp endemic to the lake, *Artemia monica* (7). In turn, *Artemia* is a crucial food source for birds along the North American Pacific Flyway (8, 9) where Mono Lake’s microbial / eukaryotic ecosystem serves a unique, multi-compartment, interlinked ecosystem role.

**Figure 1.**
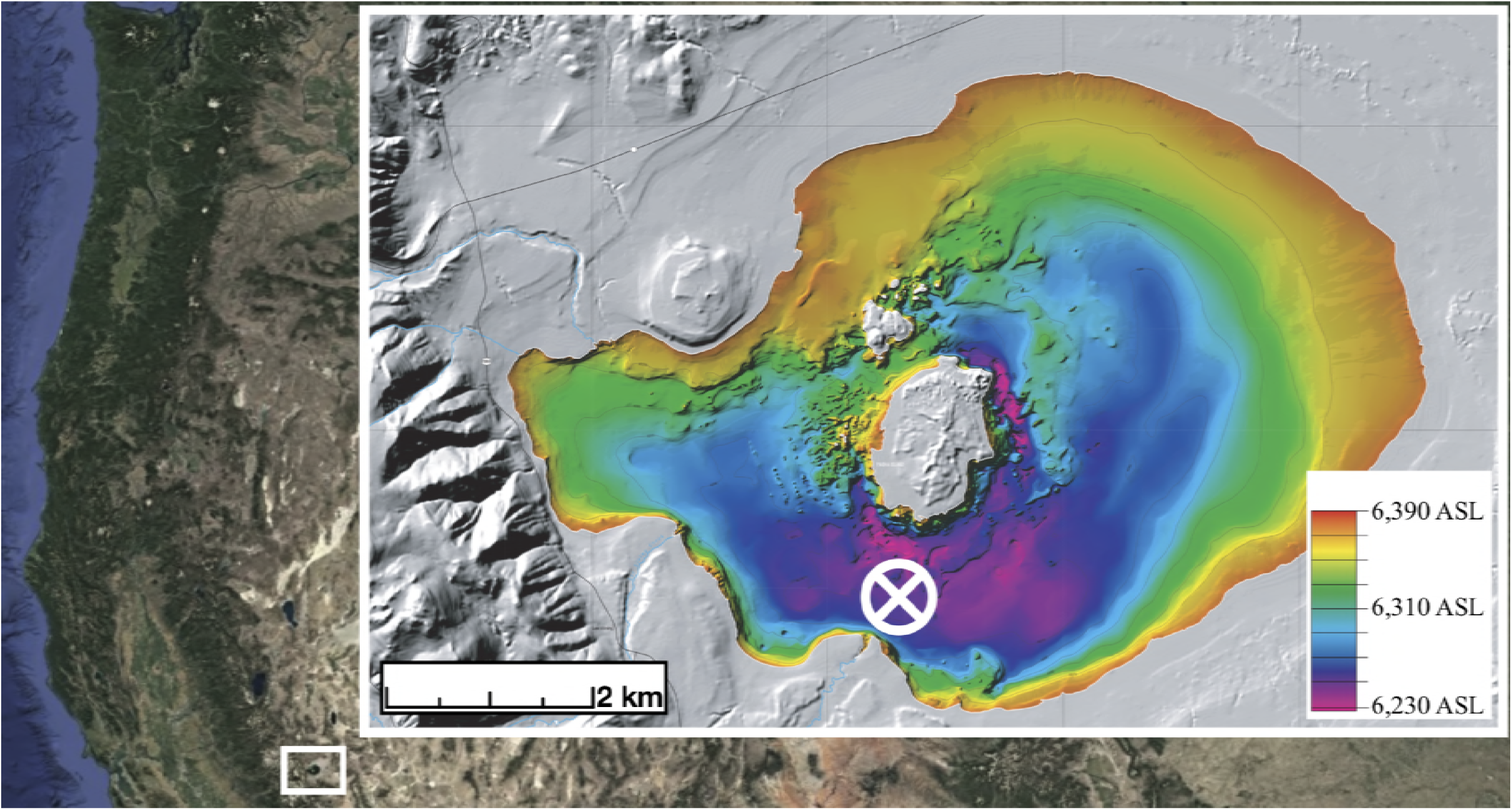
Overview of North Eastern California, with Mono Lake inset. Approximate sampling location is shown by the white circle/cross. Scale is given in kilometers. Overview image captured from Google Earth/Landsat. Inset image modified from U.S. Geological Survey Miscellaneous Field Studies Map MF-2393 (Raumann et. al 2002).

Beyond the visible macrofauna that overfly and nest at Mono Lake, the water and sediment both contain high concentrations of arsenic that have made the lake a prime location to study arsenic cycling in a natural setting (10, 11). Populations of *Gammaproteobacteria* from the family Helicobacteraceae, capable of phototrophic arsenate reduction, are commonly found within Mono Lake (12, 13). Recently, genes associated with sulfate reduction were identified below the oxycline (≈ 15 m) in Mono Lake while the lake was meromictic (i.e., stratified) (11). Prior to 2017, microbial community surveys of the lake during meromixis were only carried out using 16S rRNA gene clone library sequencing or denaturing gradient gel electrophoresis (DGGE) (14). Clone library sequencing using dideoxy chain terminator sequencing (15) is limited by sequencing coverage, and recent work using 454 Pyrosequencing (16) at Mono Lake provided additional diversity information for the lake during the onset of meromixis. However, the PCR primers chosen for previous high-throughput and clone library based amplicon surveys at Mono Lake were potentially biased (17, 18), and more recent primers for Illumina based amplicon sequencing (18) could provide a more accurate representation of lake microbial community. Furthermore, community distribution and profiling within Mono Lake during monomixis, or mixing of lake waters during a single time in a year, has yet to occur. Transcriptional profiling was recently carried out (11) with the same samples sequenced for rRNA gene analyses in another recent study (16) that describes the microbial activity from surface to below the oxycline at Mono Lake (11). A thorough description of the eukaryote responsible for much of the primary productivity in Mono Lake however, remains lacking from recent research. Such description for how, for example, the algae responsible for this primary productivity, *Picocystis strain ML,* is distributed within Mono Lake during a bloom and its impact on the ecophysiology of the lake is of crucial importance to ensure that a critical food source for migratory macrofauna is not lost.

*Picocystis* is a genus of phototrophic algae, previously characterized in other saline or alkaline environments (19, 20). *Picocystis* strain ML identified in Mono Lake (6), is a near relative of *Picocystis salinarium*, isolated from the San Francisco Salt Works in a high-salinity (∼ 85 ‰) pond (19). In addition to *P. salinarium,* other near relatives have been identified from hypersaline environments in inner Mongolia (21). *Picocystis* strain ML at Mono Lake is responsible for 100 mmol C m^−2^ d^−1^ of the primary productivity in the lake (6). Although *Picocystis* is nitrogen limited, if sufficient concentrations of ammonia are present during lake mixing and turnover (5), a bloom can occur often coinciding with periods of lake anoxia (6). It is also unknown, but possible that dissolved organic nitrogen (DON) may be a N source utilized by this eukaryote. The population density of *Picocystis* varies throughout the year, often reaching a maximum in early spring before falling as *Artemia* graze on them, reproduce and greatly reduce their number as measured by cell count (5). A possible key strategy for survival is that *Picocystis* strain ML is adapted to low-light conditions and anoxia near the bottom of Mono Lake, which prevents overgrazing by *Artemia* (22), or population decline when overgrowth reduces light transmission. Elevated concentrations of chlorophyll *a* and *Picocystis* are commonly detected below the oxycline (6). Yet, it is unknown if *Picocystis* is actively producing photosynthetic pigments and photosynthesizing under low-light conditions *in situ*, at depth. If *Picocystis* is capable of phototrophic growth below the oxycline, localized production of oxygen may disrupt localized anaerobic microbial communities in the bottom waters of Mono Lake. During a recent study, sulfate reducing microorganisms were identified alongside strictly anaerobic Clostridia at a depth of 15 m below the oxycline (11), yet conditions may not be conducive for anaerobic microbial sulfate reduction during a bloom of phototrophic algae that produce’s oxygen throughout the water column.

Mono Lake entered into a period of monomixis in 2012 corresponding to the onset of a near-record drought in the Eastern Sierra, resulting in a subsequent bloom of *Picocystis* in 2013 that failed to subside over the subsequent three years (23) and corresponded with a near record low of *Artemia* present within the lake in 2015. Lake clarity was at near-record lows and measured chlorophyll *a* concentrations were high in 2016 (23). Here, we describe the effects of an algal bloom during a period of intense drought within Mono Lake during the summer of 2016 on the distribution and abundance of the bacterial, archaeal, and eukaryotic planktonic microbial community and compare this to previously sampled years within the lake (11, 16). The possibility that the microbial community within Mono Lake could be re-populated by the sediment, groundwater, and streams that feed Mono Lake is also addressed. Finally, we determined if *Picocystis* strain ML is transcriptionally active under extremely low light levels in the lake.

## Results

### Major Ion Chemistry and Microbial rRNA Gene Copy Number within Mono Lake

At depths between 5 and 15 m, the water temperature decreased from ≈15 to ≈7 ˚C. Dissolved oxygen and photosynthetically active radiation (PAR) declined rapidly within the first 10 m, yet, fluorescence was above detectable limits throughout the sampled depths (Figure 2a). Microbial density estimated by bacterial and archaeal 16S rRNA gene copy number varied by less than 10% from 2 to 25 m. In contrast, a eukaryotic 18S rRNA gene copy number maximum was present at 20 and 25 m (Figure 2b). Major anions including sodium (Na^+^) were consistent, and near previously reported values (Table 1). Only minimal differences in anion or cation concentrations were detected within Mono Lake. Nitrate, nitrite, and sulfate were elevated at 10 m relative to 2, 20, and 25 m. No phosphate was detectable by ion chromatography (IC) from 2 to 25 m within Mono Lake, though surface water taken near shore had an average value of 0.02 mM (Table 1). Total dissolved phosphorus (potentially including phosphate and organophosphorus) measured by ICP-AES ranged from 0.59 to 0.63 mM (±0.08 mM) from surface to 25 m depth, respectively (Table 1). Most major anions and cations, and dissolved inorganic carbon, were below detectable limits in the sampled stream water and well water, with the exception of calcium which was elevated relative to Mono Lake water samples (Table 1).Individual replicate results for ICP-AES and IC are shown in supplemental table S1.

### Bacterial and Eukaryotic Microbial Community of Mono Lake, Sediment, and Surrounding Streams

After quality control a total of 694,948 DNA sequence reads were obtained, clustering into 831 operational taxonomic units (OTUs). Additional summary statistics are found in Supplementary Table S2. Chloroplast sequences were abundant across all lake water samples and were removed from further analysis. The bacterial and archaeal community differed in structure above and below the oxycline (Figure 3a). Samples taken from sediment at 10 m depth near the water sampling site also were distinct in bacterial, archaeal, and eukaryotic community structure from those in the sampled water column. Two OTUs most closely related to genera within the order Bacteroidetes decreased in relative abundance steadily with depth: *Psychroflexus* and ML602M-17, whereas unclassified Bacteroidetes remained relatively constant in abundance throughout the water column (Figure 3a). An OTU most closely related to the genus *Thioalkalivibrio* increased in abundance as depth increased. Unique to the sediment were the Euryarchaeota and the bacterial genus *Desulfonatroibacter*. An increase in the relative abundance of chloroplast sequence was noted at 20 m, increasing from 39.7 at the surface to 48.4 at 10 m, and then to 61.9 percent relative abundance at 20 m (Supplemental Figure S1). Well water taken to compare to lakewater samples contained an abundant population of OTUs most closely related to sulfur oxidizing Proteobacteria including *Thiothrix* and *Thiobacillus,* as well as Actinobacteria (*Rhodococcus*), and an abundant unclassified OTU within the Hydrogenophilaceae. Mono Lake influent stream water samples collected and examined from Rush, Mill, Lee Vining, and Wilson were distinct from samples taken from the lake itself, with the Flavobacteria, *Sediminibacterium*, and the hgcI clade of the Actinobacteria being the most abundant OTUs across all stream samples (Figure 3a). Mill was an outlier to other stream water samples, lacking abundant populations of Actinobacteria (candidatus *Planktophila*, and hgcI clade) and a lower abundance of the *Sediminibacterium* relative to Lee Vining, Rush, and Wilson streams. Community membership and distribution in the lake water column profile samples were significantly influenced (p = 0.002, R^2^= 0.90) by depth and the transition to anoxia visualized by weighted UniFrac PCoA ordination and a corresponding ADONIS test (Figure 4a).

**Table 1.** Measured geochemical parameters from the water column, as well as nearby streams and well water, representing subsurface water below Mono Lake. All values reported in millimolar (mM) as the average of triplicate samples, unless otherwise noted.

| Depth<br>(In m) | Surface | 2 | 10 | 20 | 25 | Well | Lee<br>Vining | Mill | Rush | Wilson |
| --- | --- | --- | --- | --- | --- | --- | --- | --- | --- | --- |
| Analyte |  |  |  |  |  |  |  |  |  |  |
| As<sup>a</sup> | 0.19±0.01 | 0.18±0.00 | 0.19±0.01 | 0.17±0.00 | 0.20±0.03 | BDL | BDL | BDL | BDL | BDL |
| Br<sup>a</sup> | 0.76±0.09 | 0.98±0.00 | 1.10±0.05 | 1.97<sup>d</sup> | 0.96±0.01 | BDL | BDL | BDL | BDL | BDL |
| Ca<sup>a</sup> | BDL | BDL | BDL | BDL | BDL | 3.2±3.4 | 2.7<sup>d</sup> | 1.1±1.7 | 4.5±4.0 | 1.6±2.4 |
| Cl<sup>-b</sup> | 578 ± 4.1 | 588±1.9 | 695±35 | 585±7.6 | 577±7.3 | 0.24±0.01 | 0.02±0.00 | 0.01±0.00 | 0.07±0.00 | 0.01±0.00 |
| F<sup>-b</sup> | 2.7±0.34 | 3.6±0.02 | 4.1±0.17 | 3.5±0.03 | 3.5±0.06 | 0.02±0.00 | BDL | 0.01<sup>d</sup> | BDL | BDL |
| Fe<sup>a</sup> | 0.01 | 0.01 | BDL | BDL | BDL | 0.02 | BDL | BDL | BDL | 0.01 |
| K<sup>a</sup> | 37±7.1 | 39±2.0 | 38±4.0 | 37±5.5 | 41±3.8 | 0.24±0.21 | BDL | BDL | BDL | BDL |
| Mg<sup>a</sup> | 1.9±1.4 | 1.7±1.2 | 1.0±0.02 | 1.0±0.10 | 1.4±0.30 | 2.8±4.2 | 1.5<sup>d</sup> | 0.14<sup>d</sup> | 0.34<sup>d</sup> | 5.1<sup>d</sup> |
| Na<sup>a</sup> | 1030±10 | 874±13 | 866±74 | 696±286 | 892±59 | 4.2±0.31 | 0.29±0.26 | 0.33<sup>d</sup> | 0.37±0.17 | 0.55±0.60 |
| NO<sub>2</sub><sup>b</sup> | 0.37<sup>d</sup> | BDL | 0.77<sup>d</sup> | BDL | BDL | BDL | BDL | BDL | BDL | BDL |
| NO<sub>3</sub><sup>b</sup> | 0.01<sup>d</sup> | 0.03<sup>d</sup> | 0.15<sup>d</sup> | 0.03<sup>d</sup> | 0.09<sup>d</sup> | 0.01±0.00 | 0.01±0.00 | BDL | BDL | BDL |
| P<sup>a</sup> | 0.59±0.08 | 0.62±0.05 | 0.61±0.03 | 0.59±0.04 | 0.63±0.07 | BDL | BDL | BDL | BDL | BDL |
| PO<sub>4</sub><sup>b</sup> | 0.02<sup>d</sup> | BDL | BDL | BDL | BDL | BDL | BDL | BDL | BDL | BDL |
| S<sup>a</sup> | 111±13 | 125±1.2 | 124.68±2.21 | 120.48±3.6 | 130±4.7 | 0.29±0.11 | 0.05±0.04 | 0.17±0.06 | 0.16±0.12 | 0.25±0.10 |
| SO<sub>4</sub><sup>b</sup> | 122±1.1 | 113±0.71 | 136.73±9.30 | 113.62±1.7 | 112±1.1 | 0.25±0.01 | 0.05±0.00 | 0.13±0.00 | 0.05±0.00 | 0.15±0.00 |
| DIC<sup>c</sup> | NA | 313 | 300 | 322 | 318 | 5.19 | NA | NA | NA | NA |

<sup>a</sup> Measured using ICP-AES. BDL, Below Detectible Limit.
<sup>b</sup> Measured using Ion Chromatography (IC). BDL, Below Detectible Limit.
<sup>c</sup> Dissolved inorganic carbon (DIC) reported from a single sample per site.
<sup>d</sup> Measurement from one or two samples. No standard deviation was calculated. NA indicates sample not measured.

**Figure 2.**
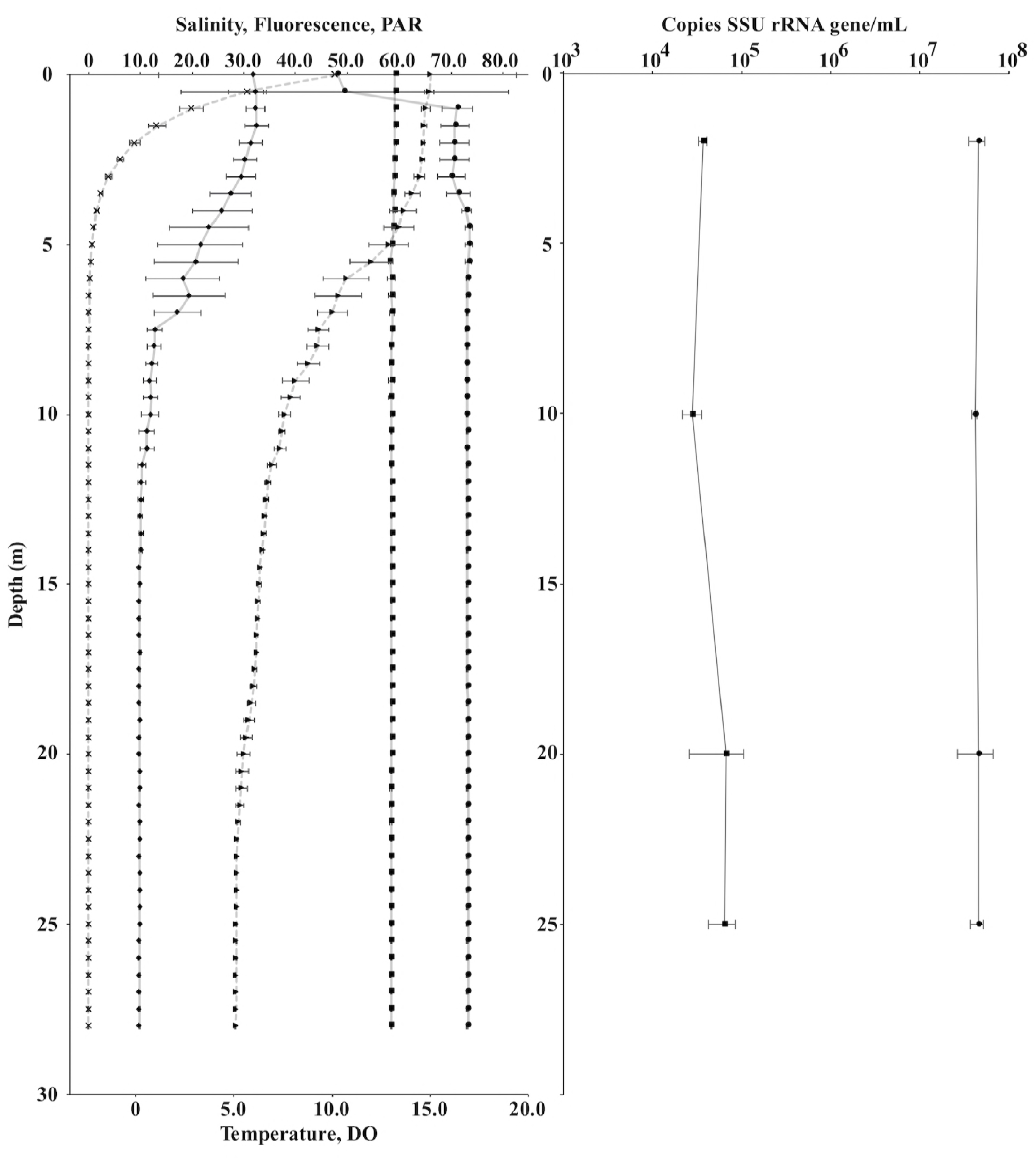
CTD measurements taken during 2016 sampling (A), with salinity (squares), fluorescence (circles), and PAR (crosses) shown on the upper axis, and temperature (triangles) and dissolved oxygen (diamonds) shown on the lower axis. Points are half-meter averages, with standard deviation shown. For clarity, lines connecting temperature and PAR are dashed. Quantification of 16S and 18S rRNA gene copy number (B) at discrete sampling depths of 2, 10, 20, and 25 m. 16S rRNA gene copy number is shown by closed circles, and 18S by closed squares, with error bars representing the mean standard deviation of triplicate biological and triplicate technical replicates.

**Figure 3.**
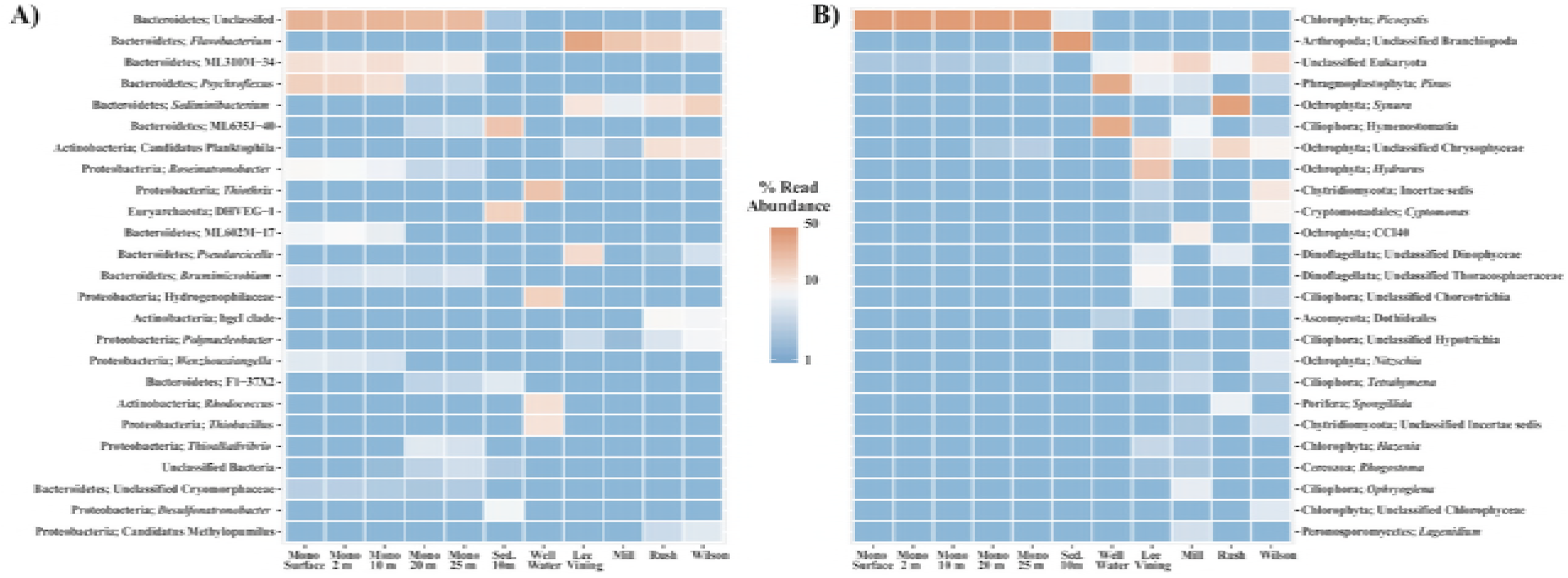
Heatmap of the top 25 OTUs within the bacteria/archaea (A) or eukarya (B). OTUs are named by Phyla, and the most likely genera.

**Figure 4.**
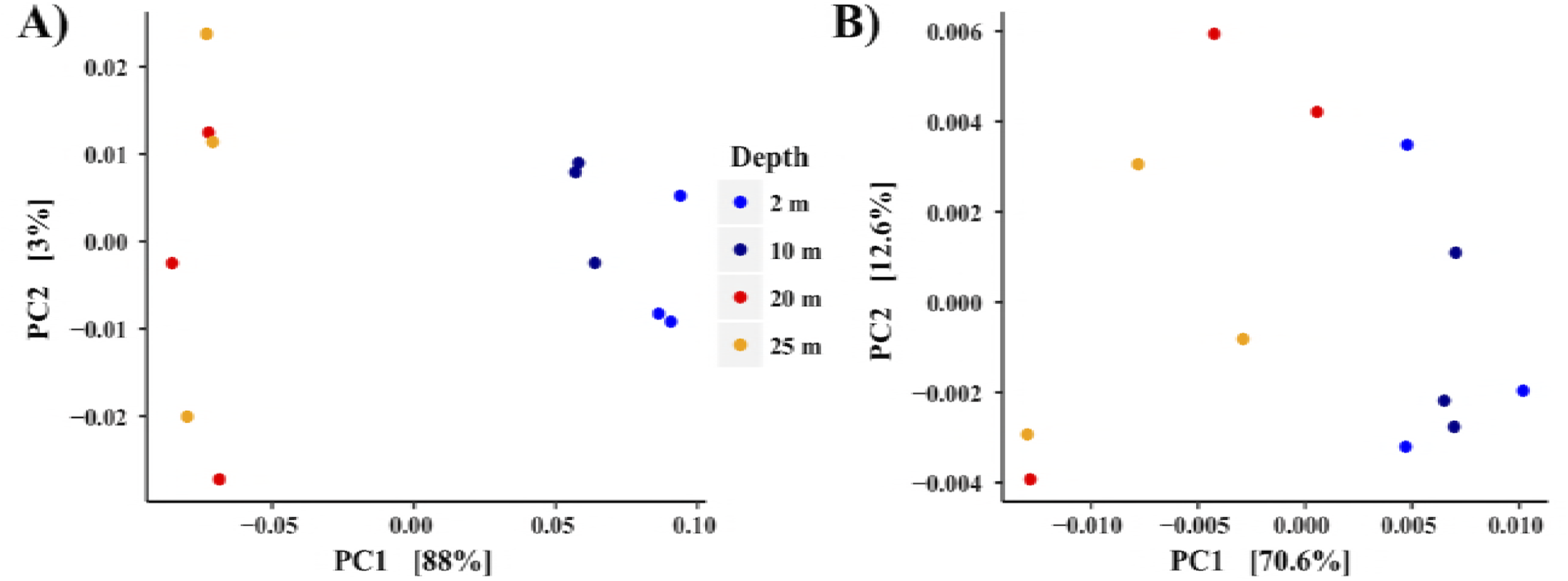
Principal component (PCoA) ordination of bacteria/archaeal (A) and eukaryotic (B) communities of water samples taken at Mono Lake. Ordination based on a weighted UniFrac distance matrix.

Compared to the observed bacterial and archaeal community, the eukaryotic community contained far fewer OTUs. Within the water column at Mono Lake, an almost homogenous distribution of OTUs most closely related to the genus *Picocystis* was observed at all depths,with a maximum of to 97.9% relative abundance at 10 m depth (Fig 3b) during the sampled bloom event of 2016. Within the sediment, an OTU of unclassified Branchiopoda was most abundant, comprising 90.9% of all sediment eukaryotic sequence. BLAST results of this OTU suggest it is most likely *Artemia monica,* endemic to Mono Lake, although because of the short sequence read length of 250 bp the identification is ambiguous. Influent stream water samples were distinct from the water and sediment of Mono Lake, with few overlapping OTUs among the samples (Fig 3b). Specifically, multiple OTUs most closely related to the Ochrophyta (Heterokont algae), Ciliophora, and Chytridiomycota were unevenly distributed across the stream and well water sampled. Community membership and distribution within the water column at Mono Lake was significantly influenced (0.017, R^2^ = 0.61) by depth and the transition to anoxia visualized by weighted UniFrac PCoA ordination and a corresponding ADONIS test, although less significantly than the bacteria and archaeal community (Figure 4b).

### Metagenomic and Transcriptomic Profiling of Mono Lake and Sediments

A summary of assembly statistics for sediment and water samples are available in Table S3. The abundance of sulfate (> 100 mM) and the lack of oxygen beg the question of whether active sulfate reduction is occurring in the dissolved organic carbon (DOC) rich waters of Mono Lake. No sulfite oxidase genes (*sox*) were identified, however genes for the complete reduction of sulfate to sulfide were identified in the sediment metagenome, and genes for reverse-dissimilatory sulfite reductases (*dsrA*) were identified in water metagenomes. No true reductive *dsrA* genes were identified in the water metagenomes. Dissimilatory sulfite reductase genes within the sediment metagenome had high (> 80 %) homology to known Deltaproteobacterial sulfate reducing microorganisms. Reductive *dsrA/B* genes were identified within the water column, identified putatively via BLAST that most closely related to known *Thioalkalivibrio dsrA/B* genes. Sulfite reductase genes did not appear to be expressed within the metatranscriptome (Table S4). Nitrate and nitrite reductases were found at 20, 25 m and within the sediment, while nitric oxide reductase (*nor*) was only identified within the sediment (Table S3). Genes associated with nitrogen fixation, including *nifH*, *D*, and *K* were found at 20 m within the water, and within the sediment metagenome. No genes associated with ammonium oxidation by bacteria or archaea (AOB/AOA) were identified. Formate-dependent nitrite reductases were identified as both genes and transcripts (Supplemental Table S4). A comprehensive list of identified transcripts is avalible as supplemental table S4 at the 10.6084/m9.figshare.6272159.

### Metagenome Assembled Genomes of Mono Lake and Sediment

After refinement binning, a metagenomic analysis identified 80 metagenome assembled genomes (MAGs) of varying completion and contamination (Supplementary Table S3). Of the 80 identified MAGs, 38 were greater than 50 percent complete, and less than 10 percent contaminated with other DNA sequence. A subset of XX of these MAGs contained rRNA gene sequence, and a putative identification was produced from these data (Supplemental figure S2). Like the rRNA gene sequencing data, MAGs indicate that microbial community composition shifted by depth and correlated to the decline in oxygen at 10 m (Figure 5). A large number of MAGs were unique to the sediment, including a Euryarchaeon (Figure 5). No archaea were found in abundance throughout the sampled water column. However, no genes associated with the production of methane were identified. Multiple MAGs were recovered from uncultivated orders within the Actinobacteria, Gammaproteobacteria, and Bacteroidetes (Table S3) including MAGs with 16S rRNA gene sequence previously identified by rRNA gene clone library sequencing at Mono Lake such as ML602J-51 (14). A summary of each genome is available in Supplementary Table S3, and figure 5. No MAGs were identified with the genes required for sulfate reduction, with only reverse-dsr genes found in MAGs. Nitrogen fixation genes (*nifH*, *D*, and *K*) were identified within 3 MAGs, two within the Gammaproteobacteria (Bin 10 and 23), as well as a single unclassified bin (Bin_11_2). One bin (Bin 45) contained photosystem II associated genes, identified within the Epsilonproteobacteria (Table S3, Figure 5). Three MAGs were identified in the EukRep filtered metagenomic sequence. A single MAG was identified with 18S rRNA gene sequence closely related to that of *Picocystis* strain ML (Supplemental Figure S2). However, this MAG appears to be contaminated with bacterial sequence, although putative searching of the identified sequence returns homology to other known algae. While the MAG should be interpreted with caution, it represents a partial genome sequence of *Picocystis* sp. from Mono Lake. The annotated genome contained no genes related to sulfur cycling, and other incomplete metabolic cycles (Supplemental Figure S3).

**Figure 5.**
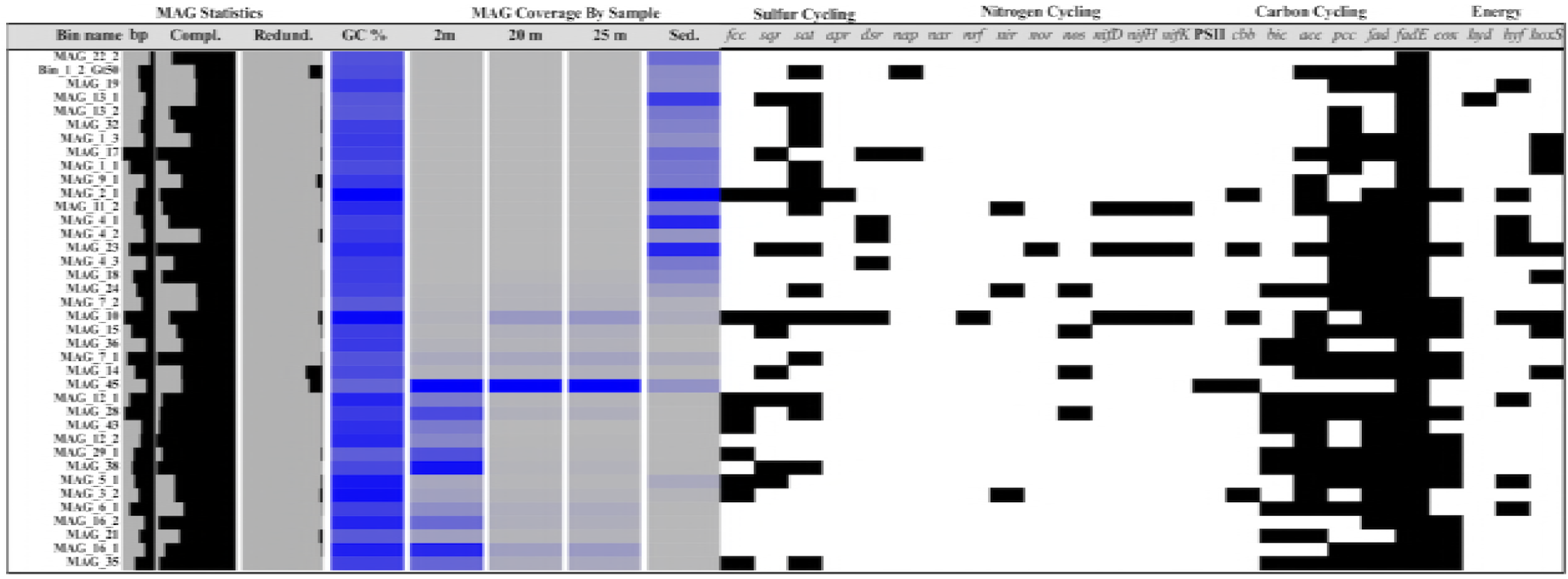
Overview of detected MAGs across sampled metagenomes and metatranscriptomes (denoted as cDNA within the figure). Color intensity from grey to blue corresponds to the coverage of each MAG within each sample. Estimates of GC content, completeness, and contamination of each MAG are also given. Presence (black) of key genes related to sulfur, nitrogen, and carbon cycling, as well as respiration are also shown. *fcc* = Sulfide dehydrogenase, *sqr* = Sulfide-quinone reductase, *sat* = sulfate adenylyltransferase, *apr* = adenosine-5-phosphosulfate reductase, *dsr* = Dissimilatory sulfite reductase, *nap* = periplasmic nitrate reductase, *nar* = nitrate reductase, nrf = nitrite reductase, *nir* = nitrite reductase, *nor* = nitric oxide reductase, *nos* = nitric oxide synthase, *nifD* = Nitrogenase molybdenum-iron protein alpha chain, *nifH* = nitrogenase iron protein 1, *nifK* = Nitrogenase molybdenum-iron protein beta chain, PSII = photosystem II, *cbb* = ribulose 1,5-bisphosphate carboxylase/oxygenase, *bic* = bicarbonate transporter, *acc* = acetyl-CoA carboxylase, *pcc* = propionyl-CoA carboxylase, *fad* = Long-chain-fatty-acid--CoA ligase, *fadE* = Acyl-coenzyme A dehydrogenase, *cox* = cytochrome c oxidase, *hyd* = Hydrogenase I, *hyf* = Hydrogenase-4, *hoxS* = bidirectional NiFe Hydrogenase.

### Metatranscriptomics Suggested Photosynthesis was Active at 25 m

Assembly of transcriptomes from 2 m and 25 m resulted in 113,202 coding sequences, and 111,709 annotated protein coding genes. No transcripts were identified with homology to known dissimilatory sulfite reductases. A total of 3,117 genes were differentially expressed (p < 0.05 false discovery rate (FDR) corrected) between 2 m and 25 m (Supplemental Table S4). More transcripts identified within the co-assembled metatranscriptome were significantly upregulated at 25 m relative to 2 m (Figure 6). Genes associated with Photosystem I and II pathways were expressed at both sampled depths (Supplemental table S4, Table 2). Expression values for photosystem I and II transcripts including *psaA/B*, *psbA/B*, and *psbC* were significantly upregulated at 25 m relative to 2 m depth (Table 2). In addition, several light-independent protochlorophyllide reductase transcripts were significantly upregulated at 25 m, while no transcripts related to chlorophyll production were significantly upregulated at 2 m (Supplementary table S4).

**Table 2.** Mean expression values of select genes identified associated with photosynthesis.

| Gene | Mean Exp. 25 m | Mean Exp. 2 m | Log<sub>2</sub> Fold Change | p value (FDR) |
| --- | --- | --- | --- | --- |
| *psaA* | 155.4 | 80.4 | 1.0 | 0.008 |
| *psaA* | 209.7 | 86.8 | 1.3 | 0.0001 |
| *psaB* | 247.0 | 123.0 | 1.0 | 0.002 |
| *psbA* | 563.7 | 176.7 | 1.7 | < 0.0001 |
| *psbB* | 241.1 | 97.4 | 1.3 | < 0.0001 |
| *psbB* | 165.4 | 74.8 | 1.1 | 0.001 |
| *psbC* | 183.5 | 84.1 | 1.1 | < 0.0001 |
| *psbY* | 23.5 | 18.4 | 0.3 | > 0.05 |
| *gyrA*<sup>a</sup> | 4.2 | 4.4 | -0.03 | - |
| *gyrB*<sup>a</sup> | 5.2 | 5.5 | -0.04 | - |

<sup>a</sup>Gyrase shown as an average expression value of all annotated *gyrA/B* transcripts at each sampled depth.

**Figure 6.**
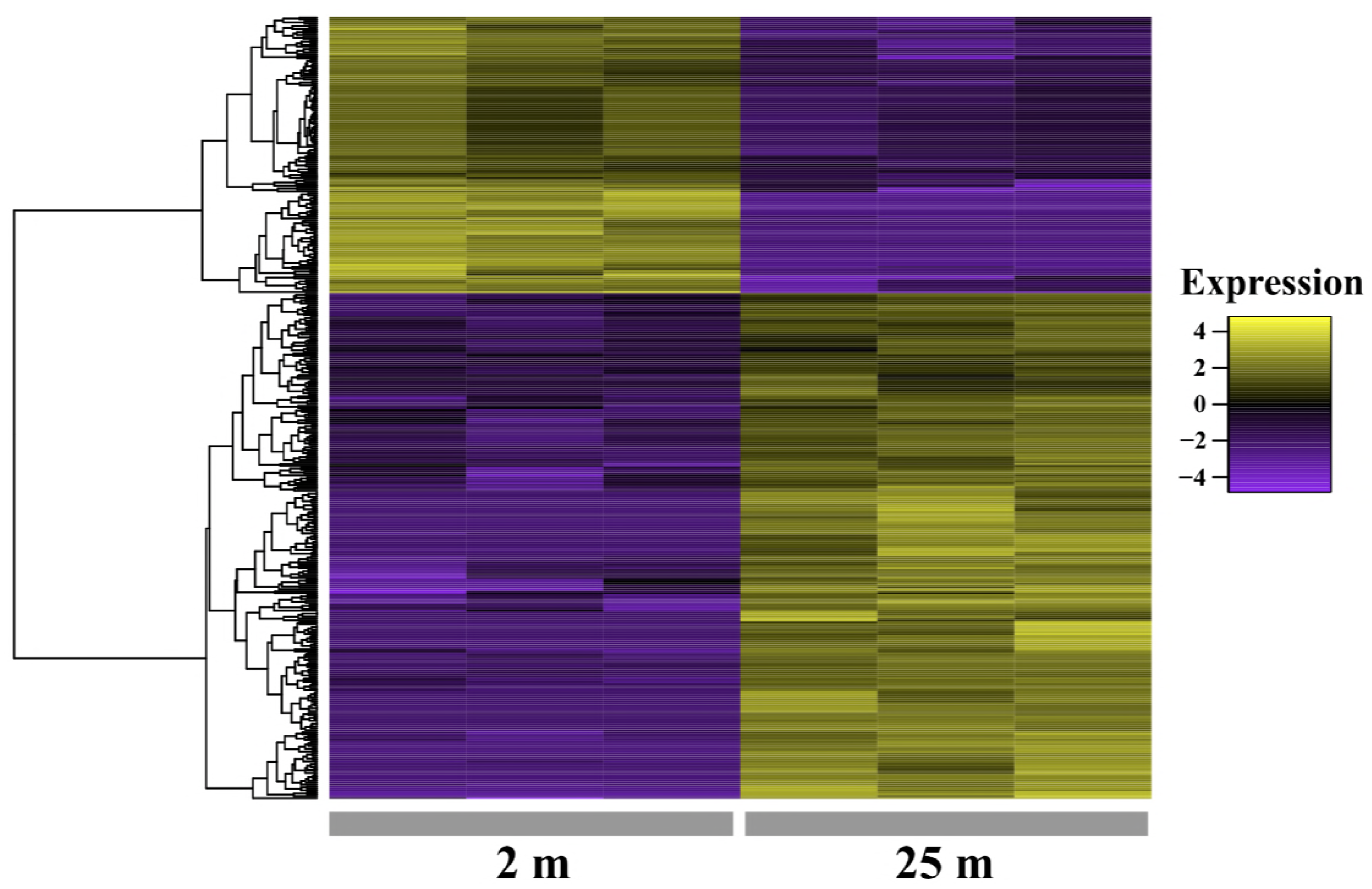
Normalized and centered expression values of *de novo* assembled transcripts significantly (FDR corrected p value < 0.05) expressed at either 2 or 25 m.

## Discussion

Beginning in late 2012, Mono Lake exhibited signs of persistent *Picocystis* blooms. Subsequently, from 2013 to 2016 both lake clarity and *Artemia* abundance declined dramatically (23). Surface concentrations of chlorophyll *a* averaged 3.8 µM in July (1994-2013), yet 2016 concentrations were ten times higher, 33.9 µM (23). The elevated chlorophyll *a* concentration and Secchi disk values (indicative of lake clarity) above 1 m suggest that Mono Lake was well within a bloom of *Picocystis*. The relative abundances of microorganisms presented here and the well-mixed major ions of Mono Lake relative to previous work (11, 14), indicated that our sampling represents the first high-throughput molecular study of Mono Lake during a *Picocystis* bloom and concurrent monomixis. Genes required for sulfate reduction to sulfide were detected only in the sequenced lake sediment, while both metagenomic and 16S rRNA gene sequencing indicated a near complete loss of the anaerobic sulfate reducing potential within the water column of Mono Lake. Instead, a mixed algal and facultatively anaerobic microbial community was present below the detectable oxycline, more similar to the near-surface microbial community than previously reported (11). It is yet unknown how the microbial community of Mono Lake will rebound after such a significant algal bloom and a decline in the population of *Artemia* within the lake.

Our survey allowed for a comprehensive evaluation of the genomic potential, and expressed genes associated with metabolic processes throughout the water column. Dissimilatory nitrate reduction to ammonium (DNRA) appeared active, with formate-dependent cytochrome c nitrite reductases detected within the transcriptome (Table S4) and formate-dependent nitrite reductase subunits within the assembled metagenomes (Table S3). No genes associated with ammonium oxidation (AOB) were identified in contrast to previous years (24) in either the transcriptome or metagenome, suggesting that the ammonia produced within the lake was assimilated, likely by the dense population of growing *Picocystis*. In addition to nitrate reduction another key anaerobic respiratory process, sulfate reduction, was largely absent from the water column.

Previous work during meromixis/non-bloom intervals has shown that sulfate reduction is a key respiratory process in Mono Lake, supporting the growth of multiple species of sulfide oxidizing aerobic microorganisms above the oxycline (11). We found that microorganisms capable of sulfate reduction were only identified in sediment metagenomic samples during the bloom.

Dissimilatory-type reverse sulfite reductases associated with sulfur oxidizing gammaproteobacterial (25) taxa were identified at 20 and 25 m, but no true reductive sulfite reductases were found in sequenced water samples. Taxa known to reduce sulfate were also only identified by 16S rRNA gene sequencing in stark contrast to previously sampled years (11, 26). Instead, the most abundant microorganisms with identifiable *dsrA/B* gene clusters were reverse-*dsr* type reductases identified previously in the Gammaproteobacterium genus *Thioalkalivibrio* (25). While lake sulfate reduction rates are typically very low (27) our data suggest a complete loss of sulfate reducing activity in the water column during a bloom. It is likely that during a bloom, sulfate reduction is repressed as more oxidizing conditions are present throughout the water column due to an increased abundance of oxygenic photosynthetic algae. Members of the Bacteroidetes were in high abundance throughout the water column, including OTUs most closely related to ML310M-34, which remained abundant through the water column and *Psychroflexus*, which decreased in abundance from 2 to 25 m as oxygen levels declined. The eukaryotic microbial community was more evenly distributed throughout the water, with *Picocystis* detected in near equivalent relative abundance throughout the water column (Figure 3b), agreeing with reported chlorophyll levels (23), as well as fluorescence values measured as a part of this study (Figure 2a).

Eukaryotic 18S rRNA gene copy number was greater at 20 and 25 m than above the oxycline by approximately 40 percent. The results were similar to previous estimates of *Picocystis* biomass during bloom events (6). *Artemia* grazing pressure was unusually low during 2016, likely allowing for the increase in *Picocystis* abundance throughout the sampled water column and accounting for the similarly low visibility (Secchi disk) readings. Additional primary productivity in the lake could also account for the oxycline shallowing from 15 m depth in 2013 (11, 14) to 10 m depth in July 2016. This expansion of anoxic waters likely limits *Artemia* populations from grazing on *Picocystis.* Lake temperature decreases at the surface relative to previous studies may also slow the metabolism of *Artemia,* resulting in reduced fecundity and increased mortality (22, 28). A decline in *Artemia* population could also impact bird mortality, though this was outside the scope of this study, and should be investigated at a later date.

A key finding of this study is the confirmation that *Picocystis* strain ML appears capable of photosynthesis under very low light conditions near the bottom of Mono Lake. Previous work suggested that *Picocystis* strain ML is capable of growth under very low light conditions, and showed elevated concentrations of chlorophyll below 15 m at Mono Lake (6). Chloroplast 16S rRNA gene sequence was most abundant at 20 m, corresponding to a peak in total 16S rRNA copy number (Figure 2b). 18S rRNA gene sequence identified as *Picocystis* were most abundant at 10 m, yet chloroplast relative abundance peaked at 20 m, near previously recorded peak depths in other recorded bloom events (6). Despite the high relative abundance of *Picocystis* throughout the water column, isolation and characterization of the *Picocystis* genome remains elusive. Binning resulted in a partial MAG with an incomplete 18S rRNA gene fragment with high similarity to the published sequence of *Picocystis* strain ML. Genome sequencing of *Picocystis,* recently isolated and sequenced twice independently (Ronald Oremland, personal communication), will allow for its genome to be removed from subsequent sequencing efforts which will simplify assembly, and enhance the resolution of bacterial and archaeal binning efforts in the future, yielding a better understanding of the microbial community responsible for the diverse metabolic potential in both the sediments and water of Mono Lake. Despite the lack of a reference genome, our transcriptomic sequencing was able to recover *Picocystis* chloroplast associated transcripts. At 25 m depth, a significant upregulation of Photosystem II was observed (Table 1, Supplemental Table S4). This, combined with the 40 percent increase in the number of 18S rRNA gene copies at 25 m relative to 2 m suggest that there is, at a minimum, a near equivalent amount of transcription of photosynthesis-associated genes throughout the water column. Recently, photosynthesis in a microbial mat was shown to be capable under extremely low light concentrations, although in a bacterial system (29). Still, the presented data suggest that under extreme low light conditions, photosynthesis may still occur. This is the first transcriptomic evidence from Mono Lake to support previous laboratory observations of *Picocystis* growing under low light conditions (6).

Our study represents the first study of Mono Lake during the height of an algal bloom and suggests significant shifts in both the bacterial and archaeal microbial community and its metabolic potential from non-bloom years (11, 16). *Picocystis* was present throughout the water column, and apparently carrying out oxygenic photosynthesis even at extremely low levels of light at depth within the lake. While Picocystis bloomed throughout Mono Lake, there was also a loss of sulfate reducing microorganisms. The lack of sulfate reduction at and below 20 m within Mono Lake is in contrast to previous work and is possibly linked to the intense drought experienced by Mono Lake from 2012 to 2016. During such a drought anaerobic microorganisms may seek refuge within the underlying sediment. By sequencing nearby sediment, we have shown that even if sulfate reduction is temporarily lost in the planktonic community of Mono Lake, the sediment may act as a “seed bank” or refugia for organisms capable of this, and likely other necessary metabolisms dependent upon overlying water / lake conditions (30).

Alternatively, the sulfate reducing microorganisms may find a better reduced substrate or fewer inhibitors in the sedimentary environment. Furthermore, the recovery of microbial populations within Mono Lake must come from its’ sediment or underlying groundwater, not from the streams that feed it as no overlapping taxa exist. Establishing if, and how, the chemistry and microbiota of Mono Lake recover after monomixis, drought, and algal bloom should be the focus of future work. Such research can be compared against our metagenomic and transcriptomic during bloom as well as previous metatranscriptomic sequencing (11) to better understand how, or if, the microbial community of Mono Lake returns to its previous state after extended periods of both monomixis and algal bloom.

## Materials and Methods

### Sampling

A vertical profile of PAR (LiCor 2π quantum sensor, 400-700 nm, E m^−2^ s^−2^), dissolved oxygen (SBE 43, mg/L^−1^), and attenuation coefficient (WetLabs transmissometer, 600 nm wavelength light source, 10 cm path length, m-1) from surface (0 m) to ∼30 m was taken using a SeaBird SBE 19 Conductivity, Temperature, and Depth (CTD) probe calibrated for use at Mono Lake. After measurements were obtained water was pumped from depth to the surface at station 6 (37.95739,−119.0316, Figure 1), sampled at 2 m, 10 m, 20 m, and 25 m the following day (due to lake conditions) using a submersible well-pump. Water was allowed to flow from the measured depth for 1 to two minutes to clear any residual water from the lines prior to sampling. *Artemia* were removed from water samples using clean cheese cloth prior to filling 1 L sterile high-density polyethylene containers. Samples were stored in a dark cooler until filtration occurred. Sediment was sampled at 10 m depth (37.9800, −119.1048) using a box-core sampling device. Well water (38.0922,−118.9919) was sampled by allowing the wellhead to flow for approximately 5 minutes before filling a 5 L HDPE container completely. For influent stream water, 1 L of water was taken from each location (Mill: 38.0230,−119.1333, Rush: 37.8883,−119.0936, Wilson: 38.0430,−119.1191) into a sterile HDPE container. Lee Vining (37.9422, - 119.1194) and was sampled with the use of a submersible pump (as above) into a sterile 1 L HDPE container.

### Geochemical Water Analysis

To characterize the water samples taken from 2 m to 25 m, major anions were measured using a Dionex ICS-90 ion chromatography system running an AS14A (4 × 250 mm) column. Major cations were also measured using a Perkin-Elmer Optima 5300 DV Inductively Coupled Plasma Optical Emission Spectrometer (ICP-OES). Both IC and ICP were conducted in the Department of Chemistry at the Colorado School of Mines. All sediment samples were extracted for ion chromatography (IC) and ICP analysis following the Florida Department of Environmental Protection method #NU-044-3.12. All fluid samples were filtered in the field using 0.22 µm PES filters. All ICP samples were acidified with trace-metal grade nitric acid as per standard procedure to ensure stabilization of all metal cations.

### Environmental Sampling, Field Preservation, and DNA/RNA Extraction of Samples

Immediately after sampling concluded, water from Mono Lake and surrounding streams were filtered onto 25 mm 0.22 µm polyether sulfone filters (Merck Milipore Corp., Billerica, MA) in triplicate. Separate triplicate filters were obtained from each water sample for DNA and RNA extraction respectively. Filter volumes are available in Supplemental Table S1. After filtration, samples were immediately suspended in 750 µL DNA/RNA shield (Zymo Research Co., Irvine, CA), and homogenized on-site using a custom designed lysis head for 1 m using a reciprocating saw. Sediment samples were immediately preserved on-site by adding sediment directly to DNA/RNA shield as above. Preserved samples were maintained on dry ice, and then stored at – 80 ˚C (RNA) or –20 ˚C (DNA) until extractions were performed. DNA extraction was carried out using the Zymo Xpedition DNA mini kit (Zymo Research Co.), and samples were eluted into a final volume of 100 µL. RNA extraction was performed using the Zymo QuickRNA Mini Prep (Zymo Research Co.) according to manufacturer’s instructions.

### rRNA Gene Sequencing Library Preparation

Libraries of bacterial, archaeal, and eukaryotic SSU rRNA gene fragments were amplified from each DNA extraction using PCR with primers (Integrated DNA Technologies Co., Coralville, IA) that spanned the ribosomal RNA gene V4 hypervariable region between position 515 and 926 (*E. coli* numbering) that produced a ∼400 bp fragment for bacteria and archaea, and a 600 bp fragment for the eukaryotes. These primers evenly represent a broad distribution of all three domains of life (18). The forward primer 515F-Y (**GTA AAA CGA CGG CCA G** <u>CCG TGYCAG CMG CCG CGG TAA</u>-3’) contains the M13 forward primer (in bold) fused to the ssuRNA gene specific forward primer (underlined) while the reverse primer 926R (5’-CCG YCA ATT YMT TTR AGT TT-3’) was unmodified from Parada et. al 2015. 5 PRIME HOT master mix (5 PRIME Inc., Gaithersburg, MD) was used for all reactions at a final volume of 50 μL. Reactions were purified using AmpureXP paramagnetic beads (Beckman Coulter Inc., Indianapolis, IN) at a final concentration of 0.8 x v/v. After purification, 4 μL of PCR product was used in a barcoding reaction, cleaned, concentrated, and pooled in equimolar amounts as previously described (31). The pooled, prepared library was then submitted for sequencing on the Illumina MiSeq platform (Illumina Inc., San Diego, CA) using V2 PE250 chemistry.

### Quantitative PCR

Total bacterial/archaeal and eukaryotic small subunit (SSU) rRNA gene count within the water column was obtained using two TaqMan based probe assays as previously described (32, 33). Briefly, both assays were carried out using 25 µL reactions containing 1x final concentration of Platinum(tm) Quantitative PCR SuperMix-UDG w/ROX (Thermo Fisher Scientific Inc.), 1.8 µM of each primer, and 225 nM of either the bacterial/archaeal, or eukaryotic probe.

### SSU rRNA Gene Analysis

Sequence reads were demultiplexed in QIIME version 1.9.1 (34), and filtered at a minimum Q score of 20 prior to clustering. Sequence reads were first denoised and then clustered into operational taxonomic units (OTUs) using UPARSE (35). After clustering, OTUs were assigned taxonomy using mothur (36) against the SILVA database (r128, (37)). Each OTU was then aligned against the SILVA r128 database using pyNAST (38), filtered to remove uninformative bases, and then a tree was created using the maximum likelihood method and the Jukes Cantor evolutionary model within FastTree 2 (39). A BIOM formatted file (40) was then produce for use in analyses downstream. To limit OTUs originating from contaminating microorganisms found in extraction and PCR reagents (41) all extraction blanks and PCR controls were processed separately and a core microbiome was computed. Any OTU found in 95% of controls was filtered from the overall dataset. Differences in community composition were estimated using the weighted UniFrac index (42). The effect of depth was tested using an adonis using the R package Vegan (43) within QIIME. Taxa heatmaps and ordination plots were generated using phyloseq (44) and AmpVis (45).

Sequencing reads for all samples are available under the project PRJNA387610. A mapping file is available both in supplemental table S2. The mapping file, as well as BIOM files used for analyses are available at 10.5281/zenodo.1247529 including an R Markdown notebook including the necessary steps to automate initial demultiplexing, quality filtering, and OTU clustering, as well as reproduce figures associated with the rRNA gene analyses.

### Metagenomic/Transcriptomic Sequencing

Metagenomic and metatranscriptomic samples were prepared using the Nextera XT library preparation protocol. Prior to library preparation, first strand cDNA synthesis was carried out using the ProtoScript cDNA synthesis kit (New England Biolabs, Ipswich, MA), followed by second strand synthesis using the NEBNext mRNA second strand synthesis module (New England Biolabs). A mixture of random hexamer and poly-A primers was using during first strand synthesis. After conversion to cDNA, samples were quantified using the QuBit HS Assay, and then prepared for DNA sequencing. Briefly, 1 ng of DNA or cDNA was used as input into the NexteraXT protocol (Illumina, Inc.) following manufacturer’s instructions. After amplification, libraries were cleaned using AmpureXP paramagnetic beads, and normalized following the NexteraXT protocol. All metagenomic and transcriptomic samples were then sequenced on the Illumina NextSeq 500 Instrument using PE150 chemistry (Illumina, Inc.).

### Metagenomic Assembly and Binning

Prior to assembly, metagenomic libraries were quality filtered and adapters removed using PEAT (46). A co-assembly was produced using MEGAHIT (47) with a minimum contig length of 5000 basepair. After assembly, quality filtered reads from individual samples were mapped to the co-assembly using Bowtie2 (44). Assembled contigs greater than 5 kb in length were first filtered to remove eukaryotic sequence using EukRep (48) and then binned into MAGs using CONCOCT (49) and refined using Anvi’o (50), in an attempt to manually reduce potential contamination or redundancy within each bin. Finally, bin quality was assessed using CheckM (51).

CheckM was also used to identify possible SSU rRNA gene fragments within each bin. Putatively identified SSU rRNA gene fragments were aligned against the SILVA 132 database (37) using SINA (52). After alignment, sequences were added to the SILVA tree by SINA, and near relatives were included to give a putative identification of MAGs containing SSU sequence. The identities of each MAG with SSU sequence are available in Supplementary Table 2.

### Metatranscriptomic Analysis

Metatranscriptome libraries were first filtered for quality and adapter removal using PEAT (46). After quality control, sequence files were concatenated into a single set of paired-end reads in FASTQ format, and then assembled *de novo* using Trinity (53). Post-assembly the Trinotate package (https://trinotate.github.io/) was used to annotate assembled transcripts. After assembly,reads were mapped against transcripts using Bowtie2 (54), and differential significance was assessed using DEseq2 (55). Assembly, annotation, mapping, and statistical analyses were carried out using XSEDE compute resources (56).

### Data Availability

Sequence data are available in the NCBI sequence read archive under the BioProject accession PRJNA387610.

## Acknowledgements

We wish to thank all participants from the International GeoBiology Course 2016 and the Agouron Institute for course funding. Ann Close and Amber Brown of USC were critical in logistics of the 2016 course and beyond. We wish to thank Tom Crowe for access to his well, and for transport on Mono Lake. Sequence data were generated by the Oklahoma Medical Research Foundation. The University of Oklahoma Supercomputing Center for Education and Research (OSCER) provided archival storage prior to sequence data submission to the NCBI SRA. This work used the Extreme Science and Engineering Discovery Environment (XSEDE), including the SDSC Comet and the TACC/IU Jetstream clusters under allocation ID TG-BIO180010, which is supported by National Science Foundation grant number ACI-1548562. A California State Parks permit to USGS and Geobiology 2016 allowed us to conduct sampling on and around Mono Lake. The funders had no role in study design, data collection and interpretation, or the decision to submit the work for publication.

## References

1. Reheis MC, Stine S, Sarna-Wojcicki AM. 2002. Drainage reversals in Mono Basin during the late Pliocene and Pleistocene. Geol Soc Am Bull 114:991–1006.

2. Cloud P, Lajoie KR. 1980. Calcite-impregnated Defluidization Structures in Littoral Sands of Mono Lake, California. Science 210:1009–1012.

3. Jellison R, Melack JM. 2003. Meromixis in hypersaline Mono Lake, California. Stratification and vertical mixing during the onset, persistence, and breakdown of meromixis. Limnol Oceanogr 38:1008–1019.

4. Nielsen LC, DePaolo DJ. 2013. Ca isotope fractionation in a high-alkalinity lake system: Mono Lake, California. Geochim Cosmochim Acta 118:276–294.

5. Melack JM, Jellison R, MacIntyre S, Hollibaugh JT. 2017. Mono Lake: Plankton Dynamics over Three Decades of Meromixis or Monomixis, pp. 325–351. In Ecology of Meromictic Lakes. Springer International Publishing, Cham.

6. Roesler CS, Culbertson CW, Etheridge SM, Goericke R, Kiene RP, Miller LG, Oremland RS. 2002. Distribution, production, and ecophysiology of Picocystis strain ML in Mono Lake, California. Limnology and Oceanography 47:440–452.

7. Warnock N. 2010. Stopping vs. staging: the difference between a hop and a jump. J Avian Biol 41:621–626.

8. Jehl JR. 1988. Biology of the Eared Grebe and Wilson’s Phalarope in the nonbreeding season: a study of adaptations to saline lakes. Studies in Avian Biology 12:1–74

9. Jehl JR. 1997. Cyclical Changes in Body Composition in the Annual Cycle and Migration of the Eared Grebe Podiceps nigricollis. J Avian Biol 28:132.

10. Oremland RS, Hoeft SE, Santini JM, Bano N, Hollibaugh RA, Hollibaugh JT. 2002. Anaerobic oxidation of arsenite in Mono Lake water and by a facultative, arsenite-oxidizing chemoautotroph, strain MLHE-1. Appl Environ Microbiol 68:4795–4802.

11. Edwardson CF, Hollibaugh JT. 2017. Metatranscriptomic analysis of prokaryotic communities active in sulfur and arsenic cycling in Mono Lake, California, USA. ISME J 54:103.

12. Hoeft SE, Blum JS, Stolz JF, Tabita FR, Witte B, King GM, Santini JM, Oremland RS. 2007. Alkalilimnicola ehrlichii sp. nov., a novel, arsenite-oxidizing haloalkaliphilic gammaproteobacterium capable of chemoautotrophic or heterotrophic growth with nitrate or oxygen as the electron acceptor. Int J Syst Evol Microbiol 57:504–512.

13. Hoeft McCann S, Boren A, Hernandez-Maldonado J, Stoneburner B, Saltikov CW, Stolz JF, Oremland RS. 2016. Arsenite as an Electron Donor for Anoxygenic Photosynthesis: Description of Three Strains of Ectothiorhodospira from Mono Lake, California and Big Soda Lake, Nevada. Life (Basel) 7:1.

14. Humayoun SB, Bano N, Hollibaugh JT. 2003. Depth distribution of microbial diversity in Mono Lake, a meromictic soda lake in California. Appl Environ Microbiol 69:1030–1042.

15. Sanger, F, Nicklen, S, and Coulson, AR (1977). DNA sequencing with chain-terminating inhibitors. Proc Natl Acad Sci U S A 74:5463–5467.

16. Edwardson CF, Hollibaugh JT. 2018. Composition and Activity of Microbial Communities along the Redox Gradient of an Alkaline, Hypersaline, Lake. Frontiers in Microbiology 9:403.

17. Klindworth A, Pruesse E, Schweer T, Peplies J, Quast C, Horn M, Glöckner FO. 2013. Evaluation of general 16S ribosomal RNA gene PCR primers for classical and next-generation sequencing-based diversity studies. Nucleic Acids Research 41:e1.

18. Parada A, Needham DM, Fuhrman JA. 2015. Every base matters: assessing small subunit rRNA primers for marine microbiomes with mock communities, time-series and global field samples. Environ Microbiol.

19. Lewin RA, Krienitz L, Goericke R, Takeda H, Hepperle D. 2000. Picocystis salinarum gen. et sp. nov. (Chlorophyta) – a new picoplanktonic green alga. Phycologia 39:560–565.

20. Krienitz L, Bock C, Kotut K, Luo W. 2012. Picocystis salinarum (Chlorophyta) in saline lakes and hot springs of East Africa. Phycologia 51:22–32.

21. Fanjing K, Qinxian J, and EJNR, 2009. Characterization of a eukaryotic picoplankton alga, strain DGN-Z1, isolated from a soda lake in Inner Mongolia, China. NREI 15:38

22. Wear RG, Haslett SJ. 1986. Effects of temperature and salinity on the biology of Artemia fransiscana Kellogg from Lake Grassmere, New Zealand. 1. Growth and mortality. J Exp Mar Bio Ecol 98:153–166.

23. Collins AG. 2017. In Response to the State Water Resources Control Board Order Nos. 98-05 and 98-07 COMPLIANCE REPORTING.

24. Ward BB, Martino DP, Diaz MC, Joye SB. 2000. Analysis of ammonia-oxidizing bacteria from hypersaline Mono Lake, California, on the basis of 16S rRNA sequences. Appl Environ Microbiol 66:2873–2881.

25. Sorokin DY, Muntyan MS, Panteleeva AN, Muyzer G. 2012. Thioalkalivibrio sulfidiphilus sp. nov., a haloalkaliphilic, sulfur-oxidizing gammaproteobacterium from alkaline habitats. Int J Syst Evol Microbiol 62:1884–1889.

26. Scholten JCM, Joye SB, Hollibaugh JT, Murrell JC. 2005. Molecular analysis of the sulfate reducing and archaeal community in a meromictic soda lake (Mono Lake, California) by targeting 16S rRNA, mcrA, apsA, and dsrAB genes. Microb Ecol 50:29–39.

27. Oremland RS, Miller LG, Whiticar MJ. 1987. Sources and flux of natural gases from Mono Lake, California. Geochimica et Cosmochimica Acta 51:2915–2929.

28. Wear RG, Haslett SJ, Alexander NL. 1986. Effects of temperature and salinity on the biology of Artemia fransiscana Kellogg from Lake Grassmere, New Zealand. 2. Maturation, fecundity, and generation times. J Exp Mar Bio Ecol 98:167–183.

29. Haas S, de Beer D, Klatt JM, Fink A, Rench RM, Hamilton TL, Meyer V, Kakuk B, Macalady JL. 2018. Low-Light Anoxygenic Photosynthesis and Fe-S-Biogeochemistry in a Microbial Mat. Frontiers in Microbiology 9:982.

30. Gibbons SM, Gilbert JA. 2015. Microbial diversity—exploration of natural ecosystems and microbiomes. Curr Opin Genet Dev 35:66–72.

31. Stamps BW, Lyles CN, Suflita JM, Masoner JR, Cozzarelli IM, Kolpin DW, Stevenson BS. 2016. Municipal Solid Waste Landfills Harbor Distinct Microbiomes. Frontiers in Microbiology 7:534.

32. Liu CM, Aziz M, Kachur S, Hsueh P-R, Huang Y-T, Keim P, Price LB. 2012. BactQuant: an enhanced broad-coverage bacterial quantitative real-time PCR assay. BMC Microbiol 12:56.

33. Liu CM, Kachur S, Dwan MG, Abraham AG, Aziz M, Hsueh P-R, Huang Y-T, Busch JD, Lamit LJ, Gehring CA, Keim P, Price LB. 2012. FungiQuant: a broad-coverage fungal quantitative real-time PCR assay. BMC Microbiol 12:255.

34. Caporaso JG, Kuczynski J, Stombaugh J, Bittinger K, Bushman FD, Costello EK, Fierer N, Peña AG, Goodrich JK, Gordon JI, Huttley GA, Kelley ST, Knights D, Koenig JE, Ley RE, Lozupone CA, McDonald D, Muegge BD, Pirrung M, Reeder J, Sevinsky JR, Turnbaugh PJ, Walters WA, Widmann J, Yatsunenko T, Zaneveld J, Knight R. 2010. QIIME allows analysis of high-throughput community sequencing data. Nature Publishing Group 7:335–336.

35. Edgar, RC (2013). UPARSE: highly accurate OTU sequences from microbial amplicon reads. Nat Methods 10, 996–998.

36. Schloss PD, Westcott SL, Ryabin T, Hall JR, Hartmann M, Hollister EB, Lesniewski RA, Oakley BB, Parks DH, Robinson CJ, Sahl JW, Stres B, Thallinger GG, Van Horn DJ, Weber CF. 2009. Introducing mothur: open-source, platform-independent, community-supported software for describing and comparing microbial communities. Appl Environ Microbiol 75:7537–7541.

37. Pruesse E, Quast C, Knittel K, Fuchs BM, Ludwig W, Peplies J, Glöckner FO. 2007. SILVA: a comprehensive online resource for quality checked and aligned ribosomal RNA sequence data compatible with ARB. Nucleic Acids Res 35:7188–7196.

38. Caporaso JG, Bittinger K, Bushman FD, DeSantis TZ, Andersen GL, Knight R. 2010. PyNAST: a flexible tool for aligning sequences to a template alignment. Bioinformatics 26:266–267.

39. Price MN, Dehal PS, Arkin AP. 2010. FastTree 2 – Approximately Maximum-Likelihood Trees for Large Alignments. Plos One 5:e9490.

40. McDonald D, Clemente JC, Kuczynski J, Rideout JR, Stombaugh J, Wendel D, Wilke A, Huse S, Hufnagle J, Meyer F, Knight R, Caporaso JG. 2012. The Biological Observation Matrix (BIOM) format or: how I learned to stop worrying and love the ome-ome. GigaScience 1:7.

41. Salter, SJ, Cox, MJ, Turek, EM, Calus, ST, Cookson, WO, Moffatt, MF, Turner P, Parkhill J, Loman NJ, Walker AW. (2014). Reagent and laboratory contamination can critically impact sequence-based microbiome analyses. BMC Biol 12.

42. Lozupone C, Knight R. 2005. UniFrac: a New Phylogenetic Method for Comparing Microbial Communities. Appl Environ Microbiol 71:8228–8235.

43. Dixon P. 2003. VEGAN, a package of R functions for community ecology. J Veg Sci 14:927–930.

44. McMurdie PJ, Holmes S. 2013. phyloseq: An R Package for Reproducible Interactive Analysis and Graphics of Microbiome Census Data. Plos One 8:e61217.

45. Albertsen M, Karst SM, Ziegler AS, Kirkegaard RH, Nielsen PH. 2015. Back to Basics – The Influence of DNA Extraction and Primer Choice on Phylogenetic Analysis of Activated Sludge Communities. Plos One 10:e0132783.

46. Li Y-L, Weng J-C, Hsiao C-C, Chou M-T, Tseng C-W, Hung J-H. 2015. PEAT: an intelligent and efficient paired-end sequencing adapter trimming algorithm. BMC Bioinformatics 16 Suppl 1:S2.

47. Li D, Liu CM, Luo R, Sadakane K, Lam TW. 2015. MEGAHIT: an ultra-fast single-node solution for large and complex metagenomics assembly via succinct de Bruijn graph. Bioinformatics 31:1674–1676.

48. West, P. T., Probst, A. J., Grigoriev, I. V., Thomas, B. C., and Banfield, J. F. (2018). Genome-reconstruction for eukaryotes from complex natural microbial communities. Genome Res 28, 569–580.

49. Alneberg J, Bjarnason BS, de Bruijn I, Schirmer M, Quick J, Ijaz UZ, Lahti L, Loman NJ, Andersson AF, Quince C. 2014. Binning metagenomic contigs by coverage and composition. Nat Meth 11:1144–1146.

50. Eren AM, Esen ÖC, Quince C, Vineis JH, Morrison HG, Sogin ML, Delmont TO. 2015. Anvi’o: an advanced analysis and visualization platform for ‘omics data. PeerJ 3:e1319.

51. Parks DH, Imelfort M, Skennerton CT, Hugenholtz P, Tyson GW. 2015. CheckM: assessing the quality of microbial genomes recovered from isolates, single cells, and metagenomes. Genome Res 25:1043–1055.

52. Pruesse E, Peplies J, Glöckner FO. 2012. SINA: accurate high-throughput multiple sequence alignment of ribosomal RNA genes. Bioinformatics 28:1823–1829.

53. Grabherr MG, Haas BJ, Yassour M, Levin JZ, Thompson DA, Amit I, Adiconis X, Fan L, Raychowdhury R, Zeng Q, Chen Z, Mauceli E, Hacohen N, Gnirke A, Rhind N, di Palma F, Birren BW, Nusbaum C, Lindblad-Toh K, Friedman N, Regev A. 2011. Full-length transcriptome assembly from RNA-Seq data without a reference genome. Nat Biotechnol 29:644–652.

54. Langmead B, Salzberg SL. 2012. Fast gapped-read alignment with Bowtie 2. Nat Methods 9:357–359.

55. Love MI, Huber W, Anders S. 2014. Moderated estimation of fold change and dispersion for RNA-seq data with DESeq2. Genome Biol 15:1–21.

56. Towns J, Cockerill T, Dahan M, Foster I, Gaither K, Grimshaw A, Hazlewood V, Lathrop S, Lifka D, Peterson GD, Roskies R, Scott JR, Wilkens-Diehr N. 2014. XSEDE: Accelerating Scientific Discovery. Computing in Science & Engineering 16:62–74.

